# Saxitoxin and tetrodotoxin bioavailability increases in future oceans

**DOI:** 10.1101/562694

**Authors:** C.C. Roggatz, N. Fletcher, D.M. Benoit, A.C. Algar, A. Doroff, B. Wright, K.C. Wollenberg Valero, J.D. Hardege

## Abstract

Increasing atmospheric levels of carbon dioxide are largely absorbed by the world’s oceans, decreasing surface water pH^1^. In combination with increasing ocean temperatures, these changes have been identified as a major sustainability threat to future marine life^2^. Interactions between marine organisms are known to depend on biomolecules, but the influence of oceanic pH on their bioavailability and functionality remains unexplored. Here we show that global change significantly impacts two ecological keystone molecules^3^ in the ocean, the paralytic toxins saxitoxin (STX) and tetrodotoxin (TTX). Increasing temperatures and declining pH increase the abundance of the toxic forms of these two neurotoxins in the water. Our geospatial global model highlights where this increased toxicity could intensify the devastating impact of harmful algal blooms on ecosystems in the future, for example through an increased incidence of paralytic shellfish poisoning (PSP). We also use these results to calculate future saxitoxin toxicity levels in Alaskan clams, *Saxidomus gigantea*, showing critical exceedance of limits safe for consumption. Our findings for TTX and STX exemplify potential consequences of changing pH and temperature on chemicals dissolved in the sea. This reveals major implications not only for ecotoxicology, but also for chemical signals mediating species interactions such as foraging, reproduction, or predation in the ocean with unexplored consequences for ecosystem stability and ecosystem services.

Climate change is not only increasing oceanic water temperatures, but also decreasing seawater pH as increasing atmospheric carbon dioxide (CO_2_) is absorbed by the ocean^1, 4^. The change occurs through the formation of carbonic acid, which further dissociates into HCO_3_^-^ and protons, leading to a predicted drop of up to 0.4 pH units by 2100, reaching mean pH levels of 7.7 in a high-emission scenario (Representative Concentration Pathway, RCP 8.5)^1^. In some coastal areas, seawater conditions of pH 7.2 are already observed temporarily^5^ and are predicted to decrease further in the future. Environmental change presents a significant challenge to marine organisms at physiological, ecological, as well as behavioural level. The current rate of pH change already impacts marine organisms’ calcification^6^, physiology and fitness^7^. Interference with animal acid-base balance and the control of neurotransmitter function have been proposed as possible mechanisms by which ocean acidification could disrupt olfactory-mediated behaviours^8^. But there is also increasing evidence that a direct impact of pH on information-carrying signalling cues and their corresponding receptors could cause info-disruption in marine chemical communication^9, 10^.

Marine organisms use a wide range of biomolecules to locate food and mating partners or to deter predators^11^. Many of these molecules also possess functional chemical groups that are sensitive to pH including hydroxyl-, carbonyl-, carboxyl-, amine-, phosphate- or sulfide-groups. Thus, changes in pH in future oceans can potentially alter a range of biological functions^2, 9^. Among these, saxitoxin and tetrodotoxin have a large variety of ecological functions at very low effective concentrations^12, 13^. They serve as antipredator defence through accumulation in cells, skin, tissue, eggs and oocytes in dinoflagellates, snails, ribbon worms, blue-ring octopus and puffer fish, or can be used as offensive weapon upon prey organisms^13, 14^. Both toxins are also released into the environment for communication purposes, e.g., as an attractant pheromone for male puffer fish^14^, or population-mediating pheromone in the harmful algal bloom (HAB) associated dinoflagellate *Alexandrium sp.*^15^. Predicted changes in climate are expected to further increase the duration, distribution and severity of HABs^16^, while ocean acidification has been directly shown to give toxic microalgae an advantage during a normal plankton bloom, resulting in their mass development and formation of HABs^17^. STX produced by *Alexandrium sp.* often accumulates in the food chain, causing paralytic shellfish poisoning (PSP) and major die-offs of fish, benthic invertebrates, and marine mammals^18, 19^, with implications for marine ecosystems as well as global food security. HABs can also cause harm to humans, including direct human mortalities, mainly due to ingesting toxic seafood, direct skin contacts with contaminated water or inhaling aerosolized biotoxins^20^.

Both neurotoxins, STX and TTX, contain functional chemical groups that are impacted by pH^21, 22^. Their more protonated forms, which are more prevalent in acidified conditions, are known to possess a more effective inhibitory capacity for ion channels^23^. Once protonated, strong electrostatic interactions of the toxins’ hydroxyl and positively charged 7,8,9-guanidinium groups with the negatively charged carboxylic side chains of the ion channels’ extracellular selectivity filter site^23, 24^ lead to a full blockage of voltage-gated sodium (NaV) channels in nerves and muscles^13^, L-type Ca^2+^ channels^23^ and voltage-gated potassium channels^13^. In comparison to their non- or partly protonated counterparts, the fully protonated forms of STX and TTX foster an even stronger electrostatic interaction with the channels, preventing ion flux, and are therefore more potent in their toxicity^23^. The longer TTX/STX is bound, the more damaging the effect on nerve and muscle fibres^23^. The effects of pH on STX and TTX toxicity have been shown in the laboratory, but have not been translated into an ecological context, nor quantified for global future ocean models. Here we calculate environmentally-mediated differences in protonation states of both TTX and STX and some of their derivatives in the context of published oceanic climate change scenarios^1^, visualise their toxicity-enhancing electrostatic differences and map their global abundance today and in future oceans.

We calculated the relative proportion of each protonation state in comparison to other states present in solution based on the p*K_a_* constants of the ionisable groups using the Henderson–Hasselbalch equation and incorporated effects of water temperature as a p*K_a_* influencing factor. The results were visualised across the pH range and compared between today’s average sea surface pH (pH 8.1), future oceanic conditions (pH 7.7)^1^ and current temporary coastal/estuarine scenarios (pH 7.2)^5^. The protonated toxic form of tetrodotoxin will increase by 9.9% under a pH change from pH 8.1 to pH 7.7 (Fig. 1a, Table 1) whilst the presence of the most toxic saxitoxin state, with protonated 1,2,3- and 7,8,9-guanidinium groups, will increase by 20.0% (Fig. 1, Table 1). Taking salt (KCl) into account alters the change towards the fully protonated form of STX to 17.0% (for details see Methods). Toxic forms of saxitoxin derivatives (neoSTX, dcSTX) also increase by 10% to 20% (see Table 1).

**Figure 1:**
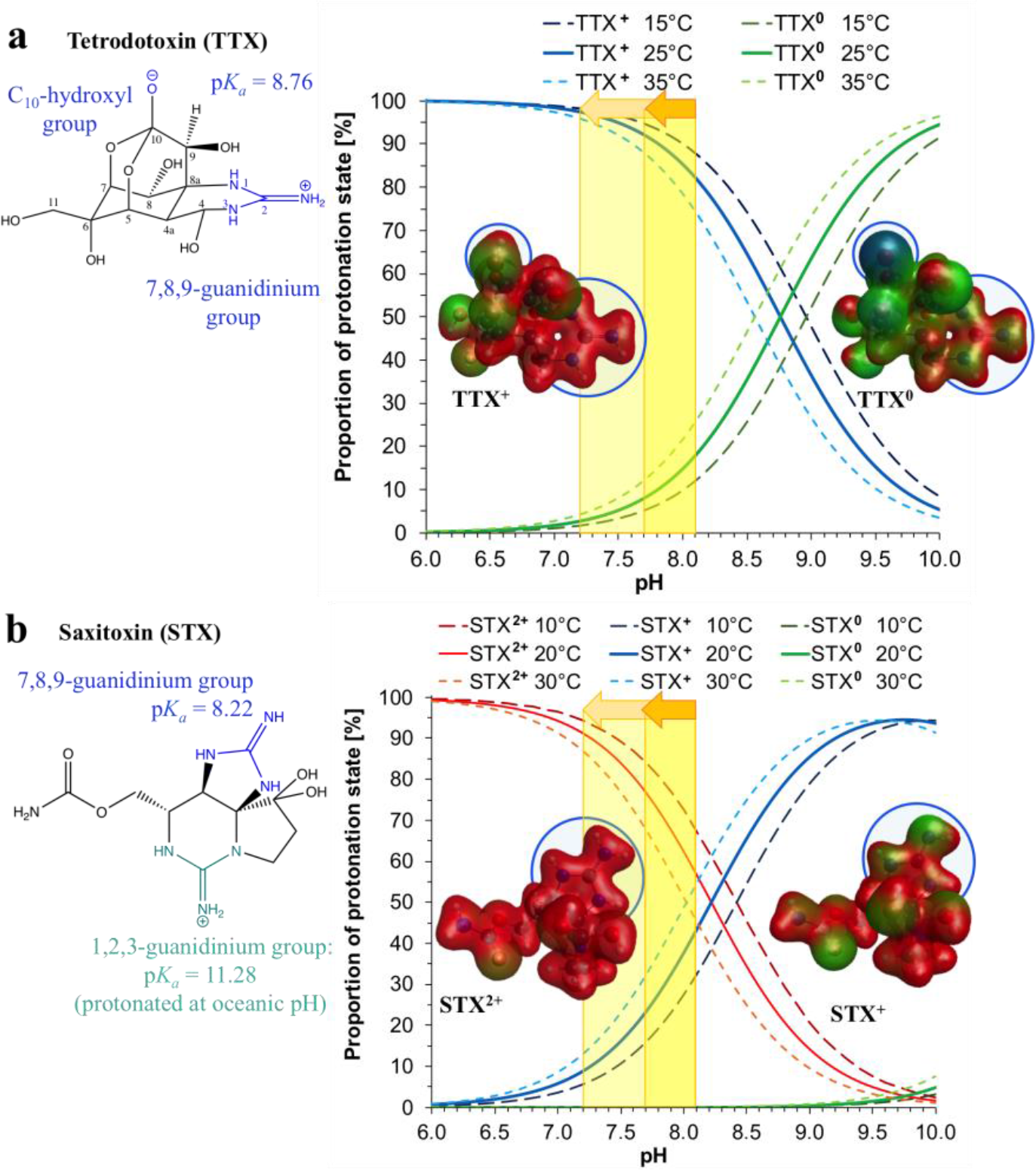
Structures, charge distribution and relative proportion of individual protonation states for (a) TTX and (b) STX. The chemical structures (left) with highlighted ionisable groups are annotated with the respective p*K_a_* values. In the proportion plots, blue (TTX) and red (STX) continuous lines represent the active toxin form abundance based on literature p*K_a_* at 25°C (TTX)/20°C (STX). The other continuous lines represent the forms with non-protonated 7,8,9-guanidinium group (green in TTX, blue in STX) and deprotonated 1,2,3-guanidinium group (green in STX). The dashed lines represent the proportions within an envelope of ±10°C around the temperature of p*K_a_* determination (+10°C - short dashes; –10°C - long dashes). The increase in toxic TTX and STX forms for the expected change in pH with ocean acidification from 8.1 to 7.7 in the year 2100 (dark yellow) and the change in pH from 8.1 to 7.2 already observed temporarily in coastal estuarine areas is highlighted by the yellow shaded areas and arrows (middle). Computationally optimised conformations (PBE0/pc-2) of the non- or partly protonated forms (TTX^0^ and STX^+^) and fully protonated toxic forms (TTX^+^ and STX^2+^) are shown with their electrostatic potential values mapped onto an electron density iso-surface. Blue indicates negative, green neutral and red positive charge.

**Table 1:** Change in abundance of TTX, STX and saxitoxin derivatives in future oceanic pH conditions.

| Compound | $pK_a$ values | Change in abundance of toxic form with pH change of 8.1 to 7.7 | Change in abundance of toxic form with pH change of 8.1 to 7.2 |
| --- | --- | --- | --- |
| TTX | 8.76 | + 9.9% ($\pm 0.3\%$/°C) | + 15.3% ($\pm 0.6\%$/°C) |
| STX | 8.22; 11.28 | + 20.0% ($\pm 0.3\%$/°C) | + 34.4% ($\pm 0.7\%$/°C) |
| STX plus 0.1M KCl | 8.39; 11.30 | + 17.0% ($\pm 0.4\%$/°C) | + 27.8% ($\pm 0.8\%$/°C) |
| dcSTX | 8.10; 10.48 | + 21.5% ($\pm 0.2\%$/°C) | + 38.8% ($\pm 0.7\%$/°C) |
| neoSTX | 6.75; 8.65; 11.65 | + 6.3% partly protonated, +5.8% fully protonated form ($\pm 0.2\%$/°C) | + 22.2% fully protonated form ($\pm 0.7\%$/°C) |

Results are based on titration data at 25°C for TTX and 20°C for STX and its derivatives. For details on $pK_a$ choice and abundance calculations see Methods.

To investigate the electrostatic properties of the protonated toxin forms hypothesised to cause enhanced toxicity, we computed lowest energy models of current and future TTX and STX protonation states using our recently developed and experimentally verified quantum chemical approach^25^. We calculated the molecular electrostatic potential (MEP) without and with the presence of water molecules (see also Supplementary Information). We visualised the charge distribution of each conformer using their MEP mapped on an electron density iso-surface to highlight molecular differences. The three-dimensional conformation of TTX and STX does not change significantly upon protonation (root mean square deviation (RMSD) of carbon atoms between protonated and non-protonated forms is ± 0.013 Å). However, the TTX^0^ and STX^+^ protonation states show a distinct charge separation while the fully protonated states TTX^+^ and STX^2+^ are overall more positively charged (Fig. 1). The most significant changes in charge from negative to positive can be observed directly at the groups subject to protonation: the oxygen bound to C-10 in TTX and the 7,8,9-guanidinium group in STX, as well as at the TTX 7,8,9-guanidinium group. Addition of explicit water molecules around the toxins is shown to have no significant impact on the charge distribution pattern (see Extended Data Figure 1). The increased positive charge at the imidazole guanidinium groups observed in our fully protonated models of both, STX and TTX, matches with the proposed mechanism of enhanced molecular toxicity^23^. The increased relative proportion of active toxin, combined with a slower degradation rate of TTX in lower pH conditions^26^ and minimal pH-effects on the receptors in the pH range of ocean acidification^27^, suggests a significantly increased bioavailability of these keystone molecules, and thus a significant increase of their toxicity, in future oceans.

To visualise the increased abundance of protonated toxic forms of STX in the ocean at a global scale we produced geospatial maps for current (Fig. 2a) and future oceanic pH conditions (Fig. 2b) based on the IPCC RCP 8.5 high-emission scenario compatible with the Paris agreement^1^. The absolute change between present and future protonated, toxic saxitoxin abundance in % is depicted in Figure 2c.

**Figure 2:**
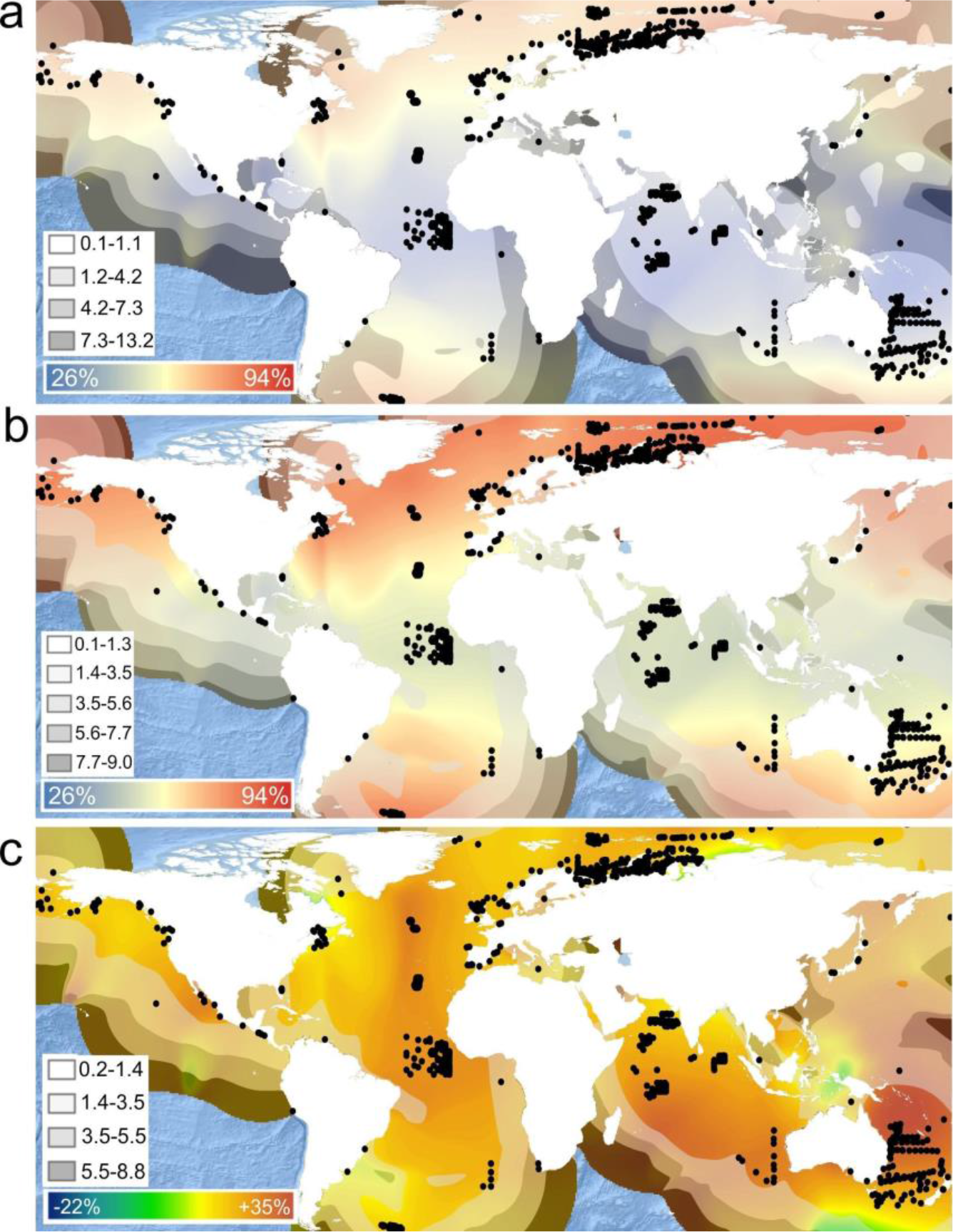
Global abundance of the toxic saxitoxin protonation state (in %) today (a), and in 2100 (b) and their difference (c). Kernel density-estimated spatial distribution maps based on concatenated records of saxitoxin-producing HABs with PSP and planktonic sampling-based occurrence records of *Gymnodinium* and *Alexandrium* dinoflagellates (black dots). a) Current mean sea surface temperature and current sea surface pH (including freshwater influx); b) under RCP8.5 estimated 2061-2100 sea surface pH and 2087-2096 predicted average sea surface temperatures; c) absolute differences in protonation state between a) and b) (zero change corresponds to light green colour). Note that b) does not incorporate estimates of freshwater influx present in a) and so might underestimate protonation state in coastal areas. The spatial distribution of the standard error of prediction (in percent, white box) is visualized as units of its standard deviation in form of grey shaded bands from transparent to dark.

The global model for present conditions shows higher levels of protonated STX, and thus greater bioavailability of the toxic form in seawater towards the poles. These increased levels of the more toxic form of STX will in the future extend towards the equator, with the Eurasian coastline of the Arctic circle reaching very high levels. The results reveal and pinpoint five “hotspots” where we predict future bioavailability of toxic STX to be significantly increased (Fig 2c): (i) the North-West Coast of the U.S.A., (ii) the Arctic Circle where *Alexandrium tamarense* blooms have already become more frequent due to warming climate^28^, (iii) the mid-Atlantic Ridge, (iv) the Indian Ocean, and lastly and perhaps most unexpectedly (v) the Coral and Solomon Seas between North-East Australia and the Solomon Islands. *Gymnodinium* and *Alexandrium* species have been recorded a little further south between Coral and Tasman Seas^29^, but any global-warming induced range shifts of these taxa^30^ could potentially lead to devastating PSP-related future HABs as indicated by current HABs around Papua New Guinea caused by the related STX-producing *Pyrodinium bahamense*^31^.

The future increase of active toxin forms shown by our geospatial interpolation models (Fig. 2) is relative to the total amount of toxin present. Combining this proportional increase in toxicity with the projected increases in HAB duration, intensity^16^ and actual higher toxin production within the cells^32^, it could result in devastating effects on marine fisheries, tourism, coastal ecosystems, and public health^20^. Recent years have seen rising numbers of STX-related PSP recordings from cold northern waters^18, 33^, such as the Barents Sea where the STX producing *Alexandrium tamarense* occurs^34^. In these areas, algal toxins, in particular STX, were identified in ten out of 13 marine stranded or harvested mammal species^33^, including humpback and bowhead whales, seals and sea otters. Since many of these affected mammals prey upon filter feeds such as the Alaskan butter clam (*Saxidomus gigantea*), a species also frequently consumed by the local people, an increase in toxicity as indicated by our maps for this region would have even more devastating direct implications.

We therefore applied our model to calculate the projected toxicity at the end of this century using current STX contents determined in butter clams collected from an affected area and found that the amount of toxic STX in butter clams from Alaska will increase in the future to levels exceeding the current US Food and Drug Administration (FDA) limit, putting marine predators and food security at risk (Fig. 3). To maintain the current recommendations for seafood safety in the future (RCP8.5 conditions), the FDA limit of 80g/100g total saxitoxin in tissue, which equals 50.4g/100g of toxic STX form (Fig. 3), will need to be reduced by over 20% to 62.8g/100g of total saxitoxin. Despite seasonal variability with clear saxitoxin summer peaks (see Extended Figure 2) all butter clam samples taken since May 2014 exceed the current FDA limit (Fig. 3b). In combination with projected increases of total STX concentration released by HABs in future ocean conditions^32^, our estimates made here for future STX toxicity may even be exceeded.

**Figure 3:**
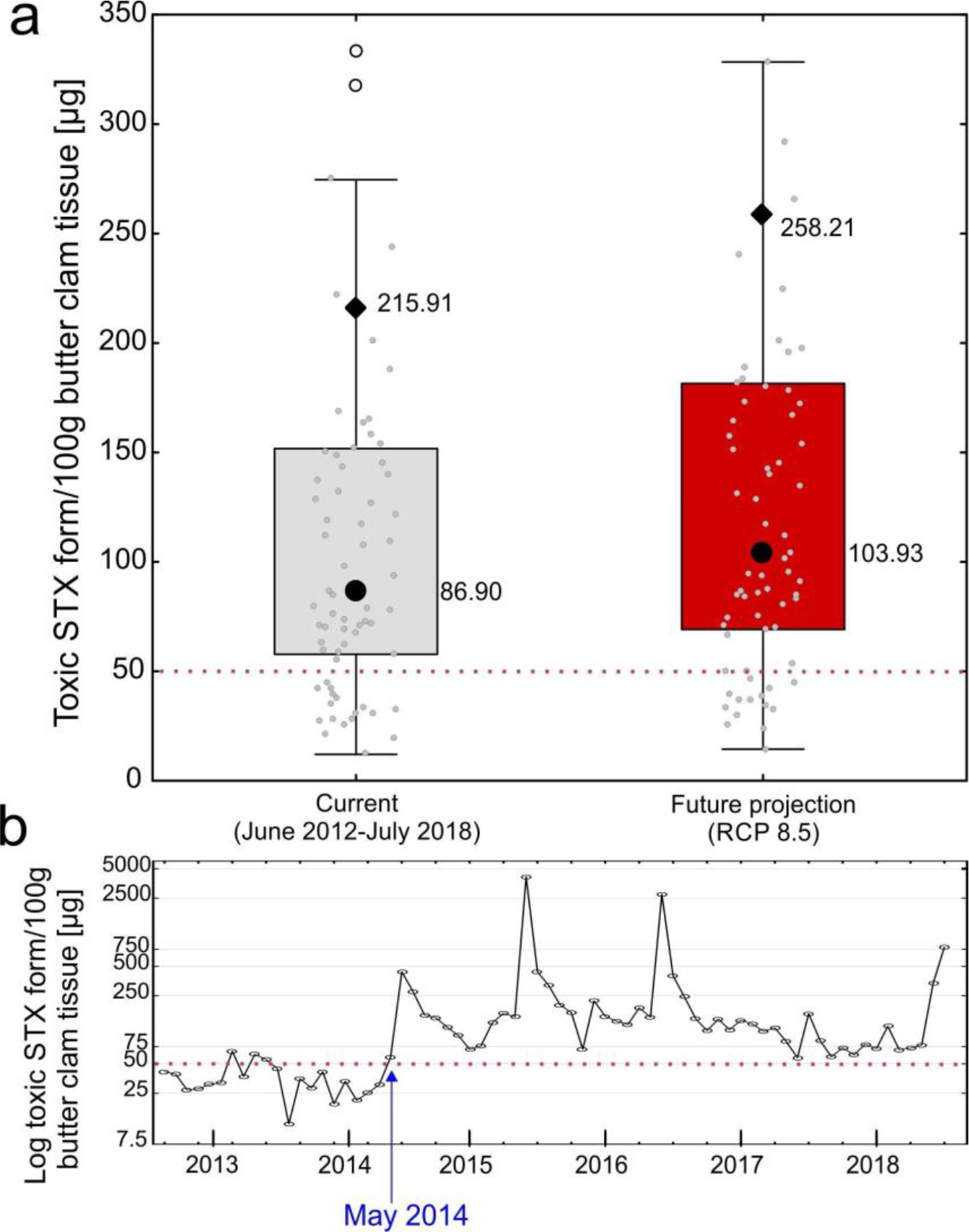
Amount of the toxic saxitoxin form present in 100g butter clam *Saxidomus gigantea* tissue at Sand Point’s Spit Beach, Alaska over the past six years (n = 70 monthly averages). Shown in a) are values (small circles) based on current and estimated future (RCP 8.5) Kernel interpolation values (see Fig. 2); the latter based on the assumption of constant amount of overall toxin and dinoflagellate abundance (values are displayed up to 350 *μ*g/100g). Large filled circles (and boxes) display the median values, diamonds are arithmetic means, and whiskers show non-outlier range. The red dashed line indicates the amount of toxic STX form (50.4 *μ*g/100g) that equals the current US Food and Drug Administration (FDA) limit of 80 *μ*g/100g total STX in seafood tissue. Additional outliers (open circles) and extreme values range to 6,580*μ*g/100g and are shown in b), which displays monthly average toxic STX contents, ranging less than the current FDA limit last in May 2014.

The most-impacted areas also encompass the Great Barrier Reef and the Solomon Islands (Fig 2c), where organisms ranging from dinoflagellates to worms, blue ring octopuses and puffer fish use STX and TTX as signalling molecules for key ecological functions^14^. Both toxins play a vital role in species interactions, such as deterring potential predators, attracting potential mates or serving as venom to overcome larger prey^14^. An imbalance of these interactions caused by altered effectiveness of these signalling molecules could significantly impact the ecological network. Many other biomolecules used by marine organisms to communicate have pH-sensitive chemical groups similar to the guanidinium groups shown here for TTX and STX^9^ and are likely to be altered by future climate change. This impact of pH can further be expected to apply not only to biomolecules but virtually any molecule dissolved in the sea that can be protonated, from marine drugs to man-made pollutants, such as pharmaceuticals, pesticides or plasticisers. However, the responses of marine organisms under future ocean conditions can be variable^8^ and difficult to predict owing to species specific differences and their potential for adaptation. A better understanding of the impacts of pH and temperature on chemicals used by marine organisms is urgently needed to assess the full risk for marine life in changing oceans.

## Acknowledgements

We acknowledge the Viper High Performance Computing facility of the University of Hull and its support team. CCR was funded by the ERC-2016-COG GEOSTICK project grant to Prof. Parsons. We thank the Quagan Tayagungin Tribe for use of their PSP Program website to access clam toxicity data. We acknowledge the World Climate Research Programme’s Working Group on Coupled Modelling and the climate modelling groups, for producing and making available their model output. For CMIP the U.S. Department of Energy’s Program for Climate Model Diagnosis and Inter-comparison provides coordinating support and led development of software infrastructure in partnership with the Global Organization for Earth System Science Portals. We would like to thank Professor D. Parsons, Energy and Environment Institute/University of Hull, and Dr. H. Bartels-Hardege, Department of Biological and Marine Sciences/ University of Hull, for valuable suggestions and discussions.

## Methods

### Calculation of protonation state abundance

Different protonation states of a molecule are present at different pH conditions. The pH at which 50% of a given ionisable group are protonated and 50% remain unchanged is given by a group-specific p*K_a_* value, which can be determined by potentiometric or NMR-based titration.^1^ For tetrodotoxin (TTX), Goto et al.^2^ obtained a p*K_a_* of 8.76 at room temperature (25°C) through multiple potentiometric titrations. For saxitoxin, Rogers & Rapoport3 found the p*K_a_* values of the ionisable 7,8,9- and 1,2,3-guanidinium groups to be 8.22 and 11.28, respectively. When performed in 1 M KCl the potentiometric titrations yielded p*K_a_* values of 8.39 and 11.30 for these groups.^3^ The p*K_a_* values of common saxitoxin derivatives were established as 8.10 and 10.48 for dcSTX by Rogers & Rapoport3 and to be 6.75, 8.65 and 11.65 for neoSTX^4^. Based on the literature p*K_a_* values, the concentration and therefore abundance of each protonation state over the pH range was calculated using the Henderson—Hasselbalch equation that relates the pH to the p*K_a_* (for details see Po & Senozan^5^ and references therein).

### Optimisation of protonation state conformers and visualisation of the charge distribution

A change in protonation states of these molecules could be accompanied by structural changes to the cues in the lowered pH of future oceans. To investigate this, we used quantum chemical calculations to obtain the energetically most favourable conformers for each possible protonation state (optimisation based on our previously published method^6^). These model conformers were then used to assess conformational differences between the protonation states, as well as differences in their molecular electrostatic potential (MEP), which describes the charge distribution around the molecule. Differences of conformers between protonation states were assessed by calculation of the root-mean square deviation (RMSD) of atom coordinates after normalisation with respect to the position of C_1_. The molecular electrostatic potential (MEP) was calculated with the GAMESS program (vJan122009R1). A three-dimensional electron density iso-surface was visualized with 100 grid points, a medium grid size and a contour value of 0.03 ea_0_^-3^ using the wxMacMolPlt program^7^ (v7.5141). The density iso-surface was coloured according to the MEP with a RGB colour scheme with red representing positive, green neutral and blue negative charge.

### Interpolation maps for spatial prediction of protonation state

In order to visualize the spatial distribution of the current protonation levels of saxitoxin as well as the effect of future changing oceanic pH and predicted increase of sea surface temperature on these, we generated Kernel interpolation maps with standard error for current and future predicted protonation states in ArcMap (V10.5.1) based on 6356 global occurrence records of saxitoxin-related PSP HABs and HAB causing dinoflagellates Gymnodinium and Alexandrium. In order to visualise future changes in estimated toxicity, we obtained raster data of current and future pH (measured as the acidity of the ocean surface), and current and future mean sea surface temperature SSH (measured as the water temperature at the ocean surface within the topmost meter of the water column in °C). Current and future STX protonation states were calculated for all 6356 point locations, which served as the data basis for the Kernel interpolation maps. More detailed methods will be uploaded upon successful publication.

### Calculation of future toxicity of STX in clam tissue and FDA limit

The saxitoxin content in *μ*g/ 100 g clam tissue was extracted from the PSP Program website of the Quagan Tayagungin Tribe^8^ for the time frame between June 2012 and July 2018 and averaged annually and for each month. The proportion of toxic STX form at current conditions as well as for future scenarios was extracted from the interpolation maps for the closest location to Spit Beach, Sand Point (Alaska). It was assumed that internal clam pH was close to the environmental pH due to the limited ability of bivalves to regulate their internal pH^9–11^. The proportional data was then used to calculate the amount of toxic STX in *μ*g 100 g clam tissue today and assuming the two future scenarios. It illustrates how the content of toxic STX in shellfish would be affected by future conditions (Fig. 3). We further calculated the amount of toxic STX currently present at the limit of 80 *μ*g/ 100 g clam tissue set by the US Food and Drug Administration (FDA)^12^, which is seen as safe to consume, and included it in Fig. 3.

